# Metabolic adjustment enhances food web stability

**DOI:** 10.1101/276550

**Authors:** Pierre Quévreux, Ulrich Brose

## Abstract

Understanding ecosystem stability is one of the greatest challenges of ecology. Over several decades, it has been shown that allometric scaling of biological rates and feeding interactions provide stability to complex food web models. Moreover, introducing adaptive responses of organisms to environmental changes (*e.g.* like adaptive foraging that enables organisms to adapt their diets depending on resources abundance) improved species persistence in food webs. Here, we introduce the concept of metabolic adjustment, *i.e.* the ability of species to slow down their metabolic rates when facing starvation and to increase it in time of plenty. We study the reactions of such a model to nutrient enrichment and the adjustment speed of metabolic rates. We found that increasing nutrient enrichment leads to a paradox of enrichment (increase in biomasses and oscillation amplitudes and ultimately extinction of species) but metabolic adjustment stabilises the system by dampening the oscillations. Metabolic adjustment also increases the average biomass of the top predator in a tri-trophic food chain. In complex food webs, metabolic adjustment has a stabilising effect as it promotes species survival by creating a large diversity of metabolic rates. However, this stabilising effect is mitigated in enriched ecosystems. Phenotypic plasticity of organisms must be considered in food web models to better understand the response of organisms to their environment. As metabolic rate is central in describing biological rates, we must pay attention to its variations to fully understand the population dynamics of natural communities.

## Introduction

Identifying the mechanisms responsible for ecosystem stability is one of the main scientific tasks in ecology (de Ruiter, 2005; Montoya et al., 2006; Rooney and McCann, 2012; Loreau and de Mazancourt, 2013). Natural ecosystems are assumed to be stable (in the sense of dynamic stability, defined by the equilibrium stability and the variability (Pimm, 1984; McCann, 2000)) thanks to many mechanisms resulting from the diversity of interacting species (MacArthur, 1955; Elton, 1958). However, mathematical models of ecosystems predicted opposite results. For instance, the theoretical study performed by May (1972) demonstrated that diversity, complexity (measured by the linkage probability between pairs of species) and the average interaction strength decreased the stability of random interaction networks (assessed by a linear stability analysis). Subsequently, many mechanisms promoting food web stability were identified and two of them inspired us to implement a new one in food web models. The first mechanism is the allometric scaling of biological rates (*e.g.* metabolic rate, feeding strength), describing them as power functions of individual body mass (Yodzis and Innes, 1992; Brown et al., 2004; Savage et al., 2004; Brose et al., 2008; Pawar et al., 2012; Kalinkat et al., 2013). These relationships provided a better prediction of species biomasses in empirical data than any other model parametrisation (Boit et al., 2012; Hudson and Reuman, 2013). In addition, allometric scaling coupled with size structured communities (*i.e.* consumers larger than their prey) lead to more stable food webs with fewer extinctions (Brose et al., 2006; Brose, 2008; Kartascheff et al., 2010).

The second mechanism is adaptive foraging. Kondoh (2003) included adaptive foraging behaviour into food web models to enable the consumers to maximise their biomass income by preferentially hunting more abundant prey. The result is dramatic, with a reversion of the pattern predicted by May: with adaptive foragers, increasing species richness and complexity enhances species persistence. Furthermore, food webs with randomly set interactions and adaptive foraging converge towards size-structured food webs with predators systematically larger than their prey (Heckmann et al., 2012). In such models, species biomasses are not the only dynamic variables, food web structures and interaction parameters are also dynamic (de Ruiter, 2005). However, one central parameter has always been considered constant in food web models: the metabolic rate. The closest examples to adjustable metabolic rates were given by Kuwamura et al. (2009), Nakazawa et al. (2011) and Wang and Jiang (2014) who considered simple models with a structured population of Daphnia including metabolically active adults and dormant eggs. In nature, however, many organisms exhibit phenotypic plasticity in the expression of metabolism (Brown et al., 2004; Glazier, 2005; Jeyasingh, 2007; Glazier, 2009a; Carey et al., 2013; Norin et al., 2015). In fact, Makarieva et al. (2005) pointed out that organisms with different body sizes can display similar metabolic rates depending on their activity. Moreover, animals from all major phyla are able to slow down their activity to face harsh conditions such as drought and starvation using body mass reduction (DeLong et al., 2014b), torpor, diapause (depression of 60-95%) or cryptobiosis (depression of 99-100%) (Guppy and Withers, 1999). Considering the metabolic activity of organisms as a constant parameter is a strong assumption despite its central role in food web models. In this study, we model the plastic response of metabolism similarly to adaptive foraging. As adaptive foraging maximises the growth rate of consumers by varying the foraging effort for the different prey, we propose that metabolic adjustment maximises the growth rate by varying the metabolic rate.

Based on prior studies on adaptive foraging, we can predict consequences of this adjustable metabolism for food web models. First, this adjustable behaviour should have a substantial impact on population dynamics. For instance, when the population density of the prey increases, consumers will raise their metabolic activity that is directly related to their consumption rate. The consequence is an increase in the predation pressure and top-down control imposed by consumers on their prey at high densities. On the contrary, in periods of starvation, consumers slow down their metabolic rate to minimise their loss in biomass caused by respiration, which keeps predator biomasses at a level high enough to avoid extinction (Chesson and Huntly, 1989; Polis et al., 1996; Chesson, 2000). In this study, we wonder whether the combination of these two effects stabilises the dynamics of the species (decreased amplitude and increased minima of population oscillations). In consequence our second prediction is that adaptive metabolic rates increase the persistence of complex food webs. As a measure of stability, we use the time variability of species biomasses (existence of fixed points and amplitude of biomass oscillation) and species persistence (proportion of surviving species in a food web).

## Material and Methods

We study the impact of metabolic adjustment on a simple tri-trophic food chain and complex food webs. Both are modified versions of the Allometric Trophic Network (ATN) (Brose et al., 2006). The complex food webs rely on the Williams and Martinez (2000) niche model for their structure and on the Yodzis and Innes (1992) predator-prey model for the dynamic equations and their parameters.

### Food web construction

The construction of the complex food webs follows the niche model (Williams and Martinez, 2000; Brose et al., 2006; Heckmann et al., 2012; Binzer et al., 2016) as it successfully predicted the food web structures of natural communities. The trophic interactions across species are set according to the algorithm detailed by Williams and Martinez (2000) with an expected connectance equal to 0.15. The basal species described by Williams and Mar-tinez (2000) are set as primary producers and the others as consumers. The niche values *n*_*i*_ (uniformly drawn in a [0, 1] interval for each of the 40 initial species) used to parametrise the niche model are also used to calculate species body mass as follows (Heckmann et al., 2012).

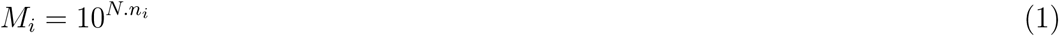

Here *N* is equal to 6, that means the biggest species is one million times larger than the smallest ones.

### Predator-prey model

The population dynamics of the food web follows the ATN model (Brose et al., 2006; Williams et al., 2007).

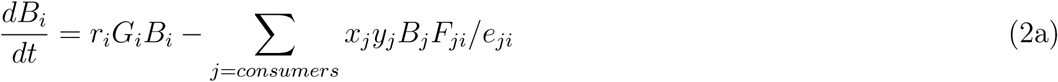

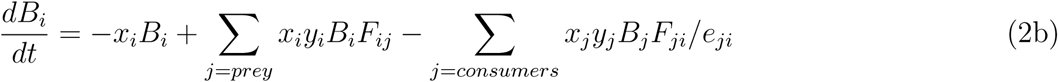

These equations describe changes in relative, biomass densities of primary producers (2a) and consumer species (2b). In these equations *B*_*i*_ is the biomass of species *i, r*_*i*_ is the mass-specific maximum growth rate of primary producers, *G*_*i*_ is the logistic growth rate of primary producers (Equation (3)), *x*_*i*_ is *i*’s mass-specific metabolic rate, *y*_*i*_ is the maximum consumption rate of consumers relative to their metabolic rate, *e*_*ji*_ is *j*’s assimilation efficiency when consuming population *i* and *F*_*ij*_ describes the realised fraction of *i*’s maximum rate of consumption achieved when consuming *j* (equation (4)). Primary producers growth rate is modelled by a logistic growth with a shared carrying capacity *K* which ensures a comparable primary production among food webs, regardless the number of primary producers (equation 3).

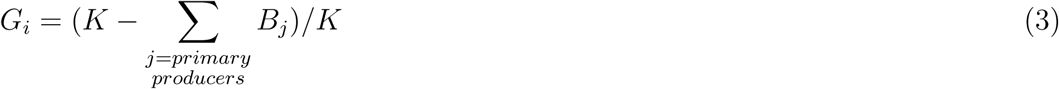

The consumption rate of prey depends on a Holling type II functional response with predator interference (Equation (4)). The preference of consumers for their prey *ω*_*ij*_ are set to 1*/p*_*i*_ with *p*_*i*_ the number of consumer *i*’s prey as we have no a priori information on preferences. Thus, all consumption rates are only driven by consumer body masses and prey biomass densities. *ω*_*ij*_ are recalculated after each extinction to follow the changes of the number of prey *p*_*i*_.

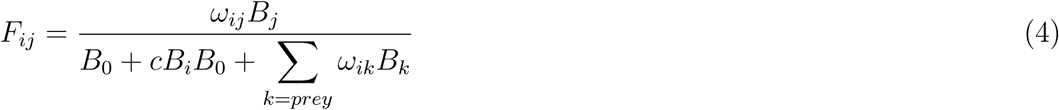

Here *B*_0_ is the half-saturation density of *i* and *c* the predator interference.

Basically, mass specific biological rates (biomass production, metabolic rate and maximum consumption rate) follow the negative-quarter power-law relationship with species body masses as described by the metabolic theory of ecology (Brown et al., 2004; Savage et al., 2004). The time scale of the system is defined by normalising the biological rates to the mass-specific growth rate of the smallest primary producer as performed by Yodzis and Innes (1992); Brose et al. (2006); Williams et al. (2007) (Equations 5a and 5b). Then the maximum consumption rates are normalised by the metabolic rates (Equations 5c). Thus, the loss due to respiration and the gain due to consumption both directly depend on the metabolic rate (Equation (2b)).

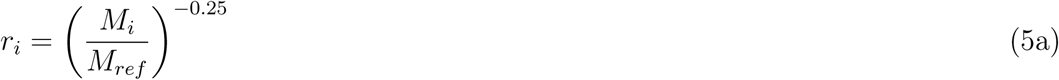

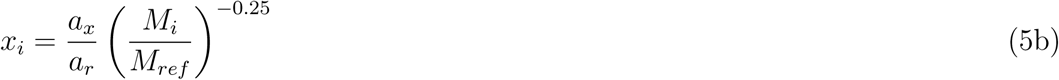

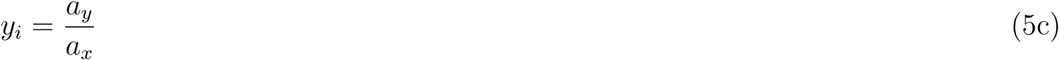

With *M* the body mass of species *i, M*_*ref*_ the body mass of the smallest primary producer, *a*_*r*_, *a*_*x*_ and *a*_*y*_ are allometric constants (see Brose et al. (2006) and Williams et al. (2007) for more details on the normalisation).

### Metabolic adjustment model

We propose to model the metabolic adjustment by an optimisation of the mass-specific net growth rate *g*_*i*_ as in adaptive foraging models (Kondoh, 2003; Uchida et al., 2007) or in body mass plasticity models (DeLong et al., 2014b). Thus, the consumer adjusts its metabolic rate to maximise the balance between ingestion and respiration that both depend on metabolic rate. Metabolic adjustment does not apply to primary producers that are considered as basal resources species with constant resource supply (Equation

(2a)).

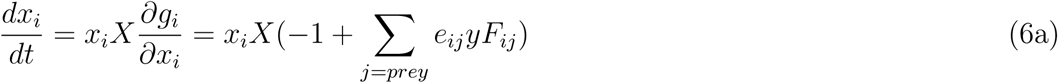

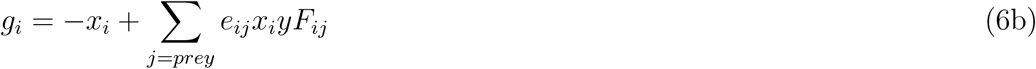

With *X* the metabolic adjustment coefficient representing the speed of the adjustment. The higher *X* is, the faster the response of species to modifications of their growth rate is. The metabolic rate is bounded by 1 and 0.001 to ensure a minimum metabolic rate and to prevent a destabilising high metabolic rate. The values predicted by the equation 5b fall in this interval that is consistent with Makarieva et al. (2005) (Supplementary material Appendix B, Fig.B5, B6,B7,B8).

### Simulations

The model is coded in *C* + + and the simulations performed with the *GSL* ODE solver. The simple tri-trophic food chain only contains a primary producer, a herbivore and a carnivore. Their body masses are respectively set to 1, 10^2^ and 10^4^. For the complex food webs, each simulation is independent from the other and only differs in the body mass distribution and the architecture of the food web. The system starts with 40 species with initial biomass of 0.1 and the metabolic rates are initialised with the values predicted by the metabolic theory of ecology (Equation 5). The simulations are performed for 10,000 time steps and only the last 1000 steps are recorded. Species persistence is the proportion of the 40 initial species that survives until the end of the simulation. Each combination of parameters is tested for 100 different food webs.

## Results

### Effect of adaptive metabolic rate on species dynamics

The first system we consider is a simple tri-trophic food chain containing a primary producer, a herbivore and a carnivore. The effects of the resource availability on species dynamics are represented by bifurcation diagrams (Fig.1). The food chain without metabolic adjustment (*X* = 0) displays large biomass oscillations whose amplitude increases with the carrying capacity *K* (Fig.1A) and the minima reaches extremely low values, especially for the herbivore (Supplementary material Appendix A, Fig.A1A). As there is no metabolic adjustment, the metabolic rates are constant (Fig.1B) and their values are those predicted by the metabolic theory of ecology (Equations 5a,b,c). The food chain with metabolic adjustment *(X* = 2) has fixed points for *K* ≤ 7 and oscillations for *K* > 7 (Fig. 1A). Despite the multi-period oscillations, the system is not chaotic (Supplementary material Appendix A, Fig.A4A). The amplitude of oscillations increases with the carrying capacity for all species but remains lower than in the food chain without metabolic adjustment. The biomass minima increases with higher values of the metabolic adjustment coefficient (Supplementary material Appendix A, Fig.A1A). The herbivore metabolic rate remains constantly at the maximum value allowed by the model, whereas the carnivore metabolic rate increases with carrying capacity *K* until it oscillates for *K* > 7 (Fig.1B).

**Figure 1:**
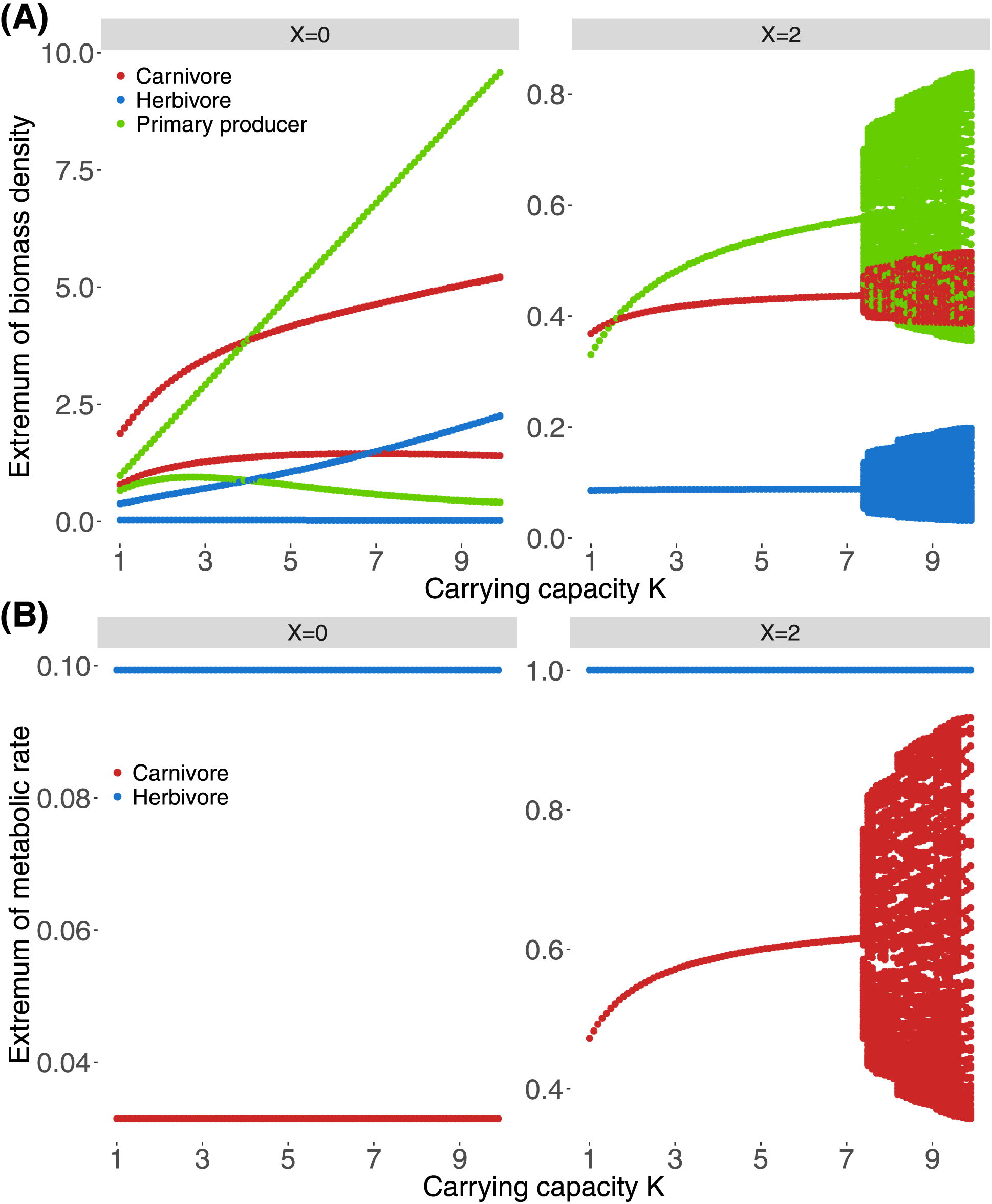
Bifurcation diagrams of the tri-trophic food-chain containing a primary producers (green), a herbivores (blue) and a carnivores (red). The bifurcation is performed along gradients in the carrying capacity *K* for **A)** biomass density and **B)** metabolic rate for a metabolic adjustment coefficient *X* = 0 or *X* = 2.

The tri-trophic food chain has fixed points along a gradient in metabolic adjustment coefficients for a carrying capacity *K* = 2 (Fig.2), except for *X* = 0 (origin of the x-axis corresponding to the situation described in Fig.1A). Increasing the metabolic adjustment coefficient increases the biomass of the herbivore and of the carnivore while it decreases the biomass of the primary producer. However, we observe an increase in the primary producer biomass and a decrease in the herbivore biomass for the low values of *X*. The metabolic rate of the herbivore is maximum for *X* > 0 and the metabolic rate of the carnivore first sharply increases with the increasing metabolic adjustment coefficient *X* and then it decreases (Fig.2B). The response is similar for *K* = 5 and *X* < 4 but for *X* ≥ 4 the system oscillates (Fig.2A), yet it is not chaotic (Supplementary material Appendix A, Fig.A4B). Increasing the metabolic adjustment coefficient does not increase the amplitude of biomass oscillations, it even decreases them for the primary producer. The biomass of the carnivore increases with *X*, the amplitude of the oscillations of its metabolic rate increases (Fig.2B) while the amplitude of its biomass oscillations remains mostly unchanged. Increasing the metabolic adjustment coefficient also increases the biomass minima of each species (Supplementary material Appendix A, Fig.A1B).

**Figure 2:**
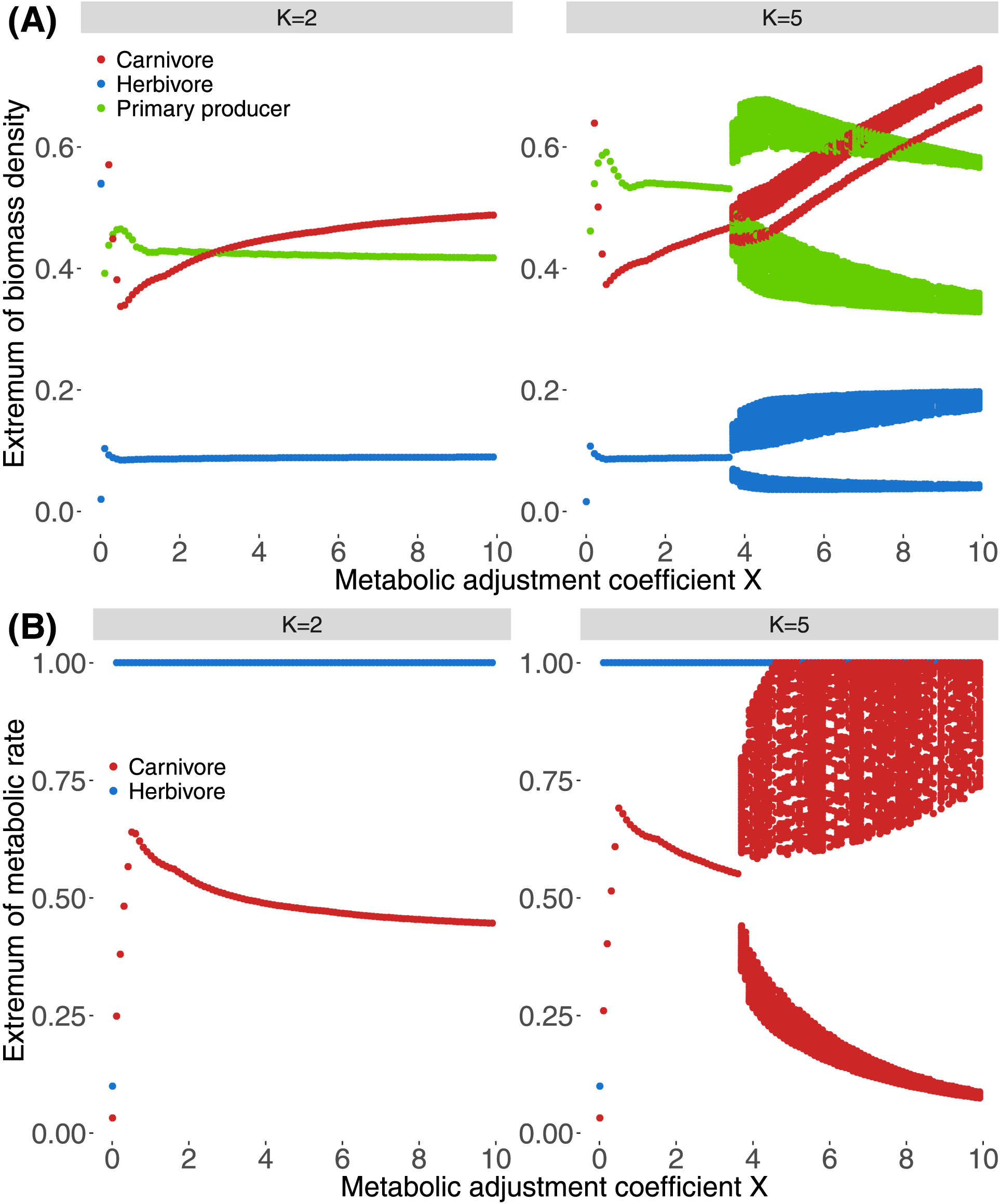
Bifurcation diagrams of the tri-trophic food-chain containing a primary producers (green), a herbivores (blue) and a carnivores (red). The bifurcation is performed along gradients in the metabolic adjustment coefficient *X* for **A)** biomass density and **B)** metabolic rate for a carrying capacity *K* = 1 or *K* = 2.

### Effect of adaptive metabolic rates on persistence

The response of stability to metabolic adjustment and enrichment in complex food webs is assessed through the average species persistence (Fig.3A). In food webs without metabolic adjustment (*X* = 0), increasing *K* does not significantly change species persistence that stays around 0.3. In food webs with metabolic adjustment (*X* > 0), for a fixed carrying capacity *K*, increasing *X* promotes species persistence, especially at low values of *K* where all species can survive. If *K* > 3, species persistence first decreases and then increases as *X* increases. For a fixed value of *X*, increasing *K* decreases species persistence and thus leads to an example of the paradox of enrichment. To sum up, enrichment through the increase of the carrying capacity has a destabilising effect on species persistence, whereas metabolic adjustment increases it substantially.

**Figure 3:**
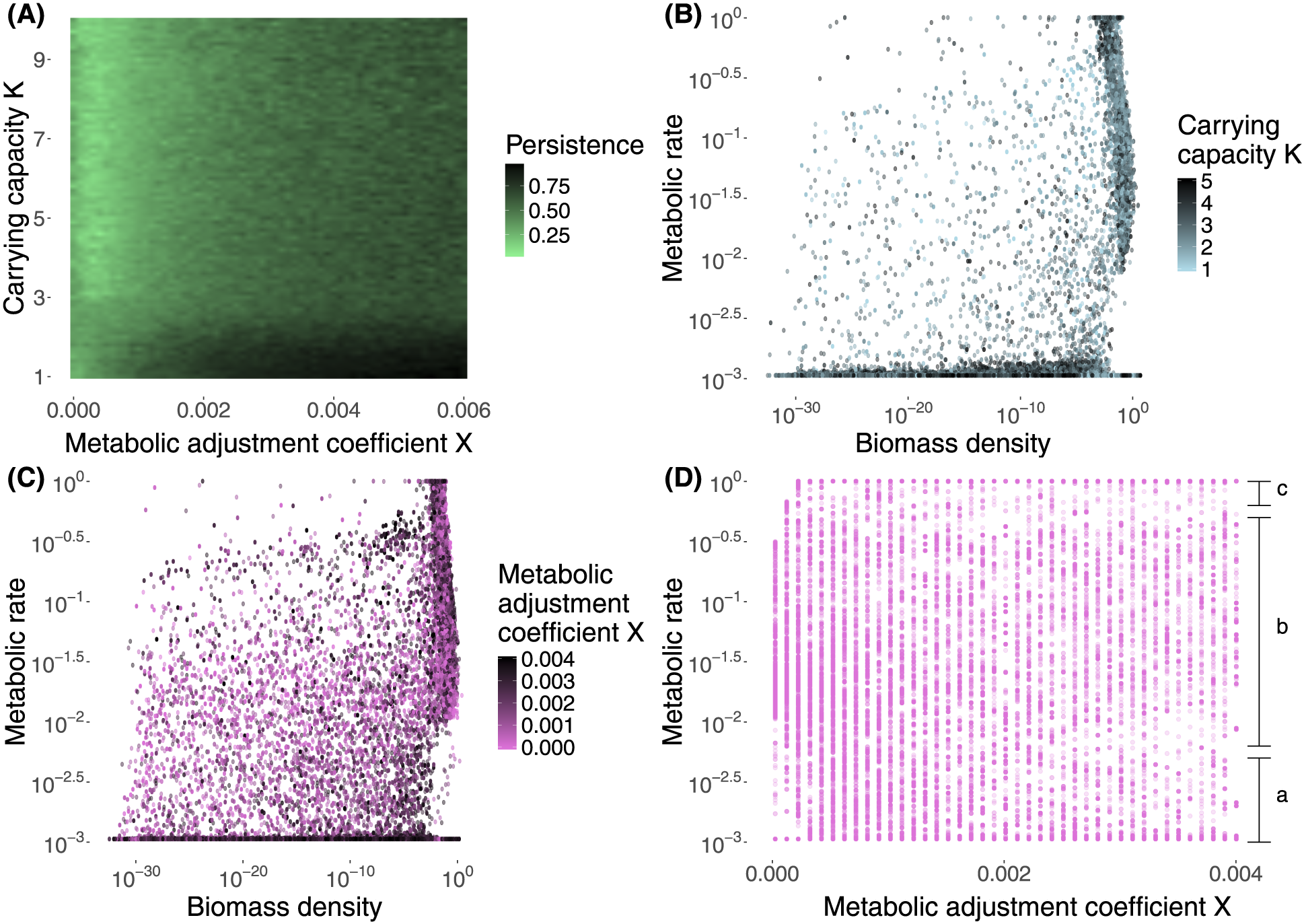
Effects of metabolic adjustment on complex food webs. **A)** Persistence of species for different values of metabolic adjustment coefficient *X* and carrying capacity *K*. Each square represent the average persistence for 100 replicates. **B)** Metabolic rate versus biomass density along gradient in carrying capacity *K* (*X* = 0.004). **C)**Metabolic rate versus biomass density along a metabolic adjustment coefficient gradient (*K* = 1.5). Each point represents one species and 100 food webs are tested for each combination of *K* and *X*. **D)** Distribution of the average metabolic rate of each species along a metabolic adjustment coefficient gradient (*K* = 1.5). The domains a, b and c represent respectively species with minimum or low metabolic rate, species with intermediate metabolic rate and species with maximum metabolic rate.

We can identify two groups of species in complex food webs: ‘slow species’ with a low biomass (*<* 10^−2^) and a low metabolic rate (*<* 10^−2^.^5^) and ‘fast species’ with a high biomass (*>* 10^−2^) and a high metabolic rate (*>* 10^22.5^) (Fig.3B and 3C). Increasing the carrying capacity *K* does not seem to change the repartition of species in these two categories (Fig.3B) while more species are in an intermediate category (low biomass and high metabolic rate) at low values of metabolic adjustment coefficient *X* (Fig.3C). *This* difference is confirmed in Fig.3D where three groups of species can be identified for *X* > 0.002: (a) species with minimum or low metabolic rate, (b) species with intermediate metabolic rate and (c) species with maximum metabolic rate. (a) species correspond to the slow species, (b) and (c) to the fast species. Such a non-differentiation of the metabolic profile of species for low metabolic adjustment coefficients may be the origin of the first decrease of species persistence with increasing *X* for *K* > 3 (Fig.3A).

## Discussion

We studied the consequences of an adaptive metabolic rate for different aspects of food web stability. We predicted that metabolic adjustment enables species to fit their metabolic rate to their energy budget and the resource availability. In times of bonanza, it allows species to increase their activity and then to exploit more resources. In harsh times, however, metabolic adjustment also lets organisms slow down their activity to save their energy until the next season of plenty (Polis et al., 1996). This behaviour is typically the case for microbial organisms that can get encysted or can produce spores (Dawes and Ribbons, 1962; Fenchel and Finlay, 1983; Glazier, 2009b) but also larger organisms that can shift between resting and activity metabolism (Glazier, 2008; Hudson et al., 2013) or hibernating (Guppy and Withers, 1999). In the case of our models, adjustable metabolic rates reduce the magnitude of biomass oscillations and increase the average biomass of carnivores. Additionally, they greatly increase the stability of complex food webs by increasing species persistence at low resource densities.

### Effect of adaptive metabolic rate on species dynamics

Our first aim was to provide a mechanistic insight in the consequences of metabolic adjustment for population dynamics. We followed prior studies employing tri-trophic food chains with allometric scaling of population parameters, which provides a fully deterministic and easily tractable system (Otto et al., 2007; Binzer et al., 2012). First, enrichment, through the increase of the carrying capacity *K*, has a destabilising effect on population dynamics (Rall et al., 2008; Schwarzmüller et al., 2015). Such a destabilisation, called paradox of enrichment, is due to the unbalance between the growth and the mortality of organisms (Rosenzweig, 1971; DeAngelis, 1992; Roy and Chattopadhyay, 2007; Rip and McCann, 2011). However, this destabilising effect is dampened by metabolic adjustment that promotes fixed points or reduces the amplitude of biomass oscillations and increases the biomass minima. Increasing the speed of adjustment (*i.e.* increasing the metabolic adjustment coefficient *X*) is destabilising because it promotes biomass oscillations, but it also increases the biomass of carnivores. We can compare our results to prior studies using adaptive foraging that inspired our modelling of metabolic adjustment (Kondoh, 2003, 2010; Křivan and Diehl, 2005; Mougi and Nishimura, 2008). The adaptability of predator attack rates or prey defences (Vos et al., 2004; Verschoor et al., 2004) also decreases in the amplitude of biomass oscillations, increases the average biomass of carnivores and keeps the minima away from the extinction threshold (Mougi and Nishimura, 2007). The outcome of these processes are similar because both rely on growth rate optimisation, which seems to highly improve the persistence of higher trophic levels that are generally most prone to extinction (Binzer et al., 2011). However, metabolic adjustment affects both the growth and the mortality rates of consumers while adaptive foraging only increases the growth rate and inducible defences decrease the mortality rate. In consequence, adaptive metabolic rates enables a better control of species dynamics, especially for top consumers whose loss rate only depends on metabolic rate and not on predation. In our tri-trophic food chain, carnivores have a highly variable metabolic rate while the herbivore’s metabolic rate always stays at the upper limit of metabolic rate range. This can be attributed to a trophic cascade: the carnivore controls the herbivore population and the primary producer thrives. Thus, the herbivore always has plenty of resources, and increasing the metabolic rate increases more the ingestion rate and the growth rate compared to the loss rate.

### Effect of adaptive metabolic rate on species persistence

Our second aim was to address the impact of an adjustable metabolic rate on the species persistence of complex food webs. The null model is a classic allometric model (Brose et al., 2006) that displays an increase in persistence with increasing carrying capacity and increase in the energy flow in the system (Dunne et al., 2005; Rall et al., 2008). As expected, adding an adjustable metabolic rate increases the species persistence at low resources levels. Similarly to the results of studies on adaptive foraging (Kondoh, 2003; Heckmann et al., 2012), higher adjustment coefficients (the metabolic adjustment in our case) increase species persistence. Such an increase in persistence can be partially attributed to the slow species with a low biomass and a low metabolic rate described in our study. However, no positive relationship between density and metabolic rate has been reported in previous studies (DeLong et al., 2014a; Yashchenko et al., 2016). Alternatively, these slow species could just be slow in getting extinct because of their very low metabolic rate (which is the loss rate in our model). However, the large diversity of metabolic rates in the fast species enables these species to better adapt to the specific situation concerning top-down control and resource availability of each food web, leading to an increased species persistence. The improvement in species persistence by the metabolic adjustment slips away as the carrying capacity increases. Our results obtained for the tri-trophic food chain demonstrate that metabolic adjustment dampens the paradox of enrichment but does not resolve it as in models with adaptive foraging (Mougi and Nishimura, 2007, 2008).

### Conclusion and perspectives

Previous models studied mechanisms similar to the metabolic adjustment by using structured populations of consumers with active adults and dormant eggs (Kuwamura et al., 2009; Nakazawa et al., 2011; Wang and Jiang, 2014). In these models, the resting eggs act as a refuge for the consumer, enabling them to escape from starvation. This mechanisms is very different of our representation of metabolic adjustment because metabolic adjustment is an energy budget optimisation process while the production of resting eggs forms a seed bank maintaining a high biodiversity (Jones and Lennon, 2010). This difference is emphasised by our divergent results. In fact, Nakazawa et al. (2011) found that the production of resting eggs leads to more stable population dynamics as it responds more to seasonality than to non-seasonal variation in resource availability (in this case the effect of resting eggs is weak). Metabolic adjustment (*i.e.* response to resource availability) in food webs deeply changes the outcome of the model. In fact, adjustable metabolic rates greatly increase stability regarding many criteria: they increase the average biomass of top trophic levels, decrease the variability in population biomass density and increase the minima of population biomass density, keeping them away from the extinction threshold. Including metabolic adjustment in food web models improves the representation of the diversity of organisms whose metabolic activity is not predicted by the metabolic theory of ecology (Guppy and Withers, 1999; Glazier, 2005; Makarieva et al., 2008; DeLong et al., 2014b). More broadly, considering phenotypic plasticity (as it was extensively done for adaptive foraging or inducible defences for instance) is crucial to better understand the fast response of organisms to environmental changes (Marshall and McQuaid, 2011; Marshall et al., 2011; Magozzi and Calosi, 2015). Interesting future directions in this research agenda would be to extend metabolic adjustment to primary producers depending on the supply of non-biotic resources affected by seasonality (*e.g.* nutrients, sun light, water…) or to include more parameters such as the attack rate in the list of biological rates directly affected by the adjustable metabolic rate. Finally, it would also be interesting to set the metabolic adjustment coefficient *X* as an allometric parameter because single cell organisms are expected to respond faster than large animals for instance. Overall, adjustable metabolic rates holds great potential to represent the biology of many species in natural communities as metabolic rate plays a central role in describing species biological functions.

## Acknowledgments

We would like to thank Christoph Digel and Björn Rall for their help during this study. I would also like to thank members of the journal club of the iEES and Marie-Hélène Berthet for their helpful review. We thank the École Normale Supérieure and the PhD program “Ecole Doctorale Frontières du Vivant (FdV) – Programme Bettencourt” for their financial support. U.B. acknowledges support by the German Research Foundation (FZT 118).

## Data accessibility

All data are included in the manuscript and its supporting information. The codes are available on Zenodo and GitHub (doi:10.5281/zenodo.1170138).

## Supplementary materials

Supplementary material (available online as Appendix XXXXX (insert manuscript number) at LÄNK). Appendix 1–2

---

**Table 1:** Parameters and variables used in the model

| Variable | Value | Description |
| --- | --- | --- |
| $B_i$ | $\text{kg.m}^{-2}$ | biomass density of species $i$ |
| $r_i$ | dimensionless | scaled mass specific maximum growth rate of species $i$ |
| $x_i$ | dimensionless | scaled mass specific metabolic rate of species $i$ |
| $y_i$ | 8 | scaled mass specific maximum consumption rate |
| $e_{ji}$ | 0.45 | assimilation efficiency of species $i$ by species $j$ (herbivores) |
| | 0.85 | assimilation efficiency of species $i$ by species $j$ (carnivores) |
| $G_i$ | dimensionless | density dependent growth rate of species $i$ |
| $F_{ij}$ | dimensionless | functional response of species $i$ feeding on species $j$ |
| $B_0$ | $0.5 \text{ kg.m}^{-2}$ | half saturation density for consumer functional response |
| $c$ | $0.5 \text{ m}^2.\text{kg}^{-1}$ | predator interference |
| $\omega_{ij}$ | 1/nbr prey | predator $i$ preference for species $j$ |
| $a_x/a_r$ | 0.138 | metabolic rate allometric constant (primary producers) |
|  | 0.314 | metabolic rate allometric constant (invertebrates consumers) |
| $X$ | dimensionless | metabolic adjustment coefficient |
| $K$ | $\text{kg.m}^{-2}$ | carrying capacity of primary producers |

Note: All these parameters come from Brose et al. (2006).

